# ModiBodies: A computational method for modifying nanobodies to improve their antigen binding affinity and specificity

**DOI:** 10.1101/820373

**Authors:** Aysima Hacisuleyman, Burak Erman

## Abstract

Nanobodies are special derivatives of antibodies, which consist of only a single chain. Their hydrophilic side prevents them from having the solubility and aggregation problems of conventional antibodies, and they retain the similar size and affinity of the binding area to the antigen. Nanobodies have become of considerable interest for next-generation biotechnological tools for antigen recognition. They can be easily engineered due to their high stability and compact size. They have three complementarity determining regions, CDRs, which are enlarged to provide a similar binding surface to that of regular antibodies. The binding residues are more exposed to the environment. One common strategy to improve protein solubility is to replace hydrophobic residues with hydrophilic ones on the binding surface which contributes to both stability and solubility of nanobodies.[1] Here, we propose an algorithm that uses the 3D structures of protein-nanobody complexes as the initial structures and by successive mutations in the CDR domains to find optimum binding amino acids for hypervariable residues of CDRs to increase the binding affinity and nanobody selectivity. We used the MDM4-VH9 complex, (PDB id 2VYR), fructose-bisphosphate aldolase from *Trypanosoma congolense*, (PDB id 5O0W), and human lysozyme, (PDB id 4I0C). as benchmark studies and identified similar amino acid patterns in hypervariable residues of CDRs with experimentally optimized ones. According to this method, better binding nanobodies can be generated by using this algorithm in a short time. We suggest that this method can complement existing immune and synthetic library-based methods, without the need of experiments or large libraries.

## Introduction

Antibodies, also called immunoglobulins, are Y shaped proteins that are key elements in the adaptive immune system. They recognize unique parts of foreign targets, antigens, in the blood or mucosa and inactivate them. They activate complementary systems to destroy bacterial cells and facilitate phagocytosis of foreign substances. They are composed of two identical copies of heavy (approximately 50 kDa) and two identical copies of light chains (approximately 25 kDa). Heavy chains are connected to each other by a disulfide bond and each heavy chain is connected to a light chain by another disulfide bond. Each antibody producing B-cell produces a unique type of antibody. There are 5 antibody isotypes: IgG, IgM, IgD, IgE and IgA all differ in their heavy chains. Variance in their heavy chains result in them to bind different antigens.[2] Antibody derived biologics have produced impressive therapeutic results. Advances in antibody engineering provided improved innovative molecules with impressive achievements in treatments of several hematological malignancies and tumors. The advantages of them are their enhanced effector function with reduced immunogenicity with prolonged half-life with reduced side effects, but their large size and hydrophobic binding surfaces pose obstacles in their stability and access to the antigen. Due to these limiting factors, new antibody-formatted treatment options with similar binding specificity and higher stability are required.

Nanobodies, special derivatives of antibodies, that consist of only heavy chains are produced by members of Camelidae family (*Llama glama*, *Vicugna pacos*, *Camelus dromedaries, Camelidae, Camelus bactrianus*), nurse sharks, *Ginglymostoma cirratum*, wobbegongs, *Orectolobus* and spotted ratfish, *Hydrolagus colliei*. They are highly specific and exert high affinity towards their target. They can be easily expressed in microorganisms. Their toxicity is low and and their tissue penetration is not limited due to their small size. The regions on nanobodies that are responsible for binding and recognition are called complementarity-determining regions (CDR).[3] The antigen binding region, paratope, diversity of antibodies are derived from their heavy and light chain combinations. Nanobodies are only composed of heavy chains compensate this diversification by taking many more loop architectures without any structural restrictions in their CDR’s. Due to their biochemical functionality and economic benefits, interest in nanobodies has grown in biotechnology and medicine. In order to use nanobodies as a therapeutic reagent, their key properties such as their stability, binding specificity and affinity to the target antigen must be optimized.

The target specificity of a nanobody is provided by its CDRs. Nanobodies comprise of three CDR loops; CDR1, CDR2 and CDR3. CDR3 is the dominating contributor in antigen recognition while the impact of the first and the second CDR loops are limited. Residue numbers of the CDR amino acids can vary but a CDR loop can be recognized by the presence of certain amino acid repeats. These repeats are composed of evolutionarily fixed and variable regions. The template of the fixed and variable residues differs within species. There is a statistical study conducted on known, stable llama derived nanobodies, showing which CDR residues are conserved and which CDR residues are hypervariable.[4] The increased frequency of some residues in the hypervariable positions supports convergent evolution in those regions.

The structure, consensus sequence and the hypervariable positions for *Llama glama* derived nanobodies are given in *Fig 1.*The frequency scheme of the highly variable regions shown with asterisks (*) in *Fig1b* are given as; 14%-Y, 12%-G, 10%-S, 9%-D, 7%-T, 6%-R, 6%-A, 5%-L, 5%-V, 4%-N, 3%-F, 3%-E, 3%-I, 3%-W, 2%-Q, 2%-K, 2%-H, 0%-C and 0%-M. In nature, antibodies are mutated naturally to increase the binding affinity and the specificity towards their target antigen. The hypervariable residues in CDRs determine the specificity of a nanobody. Similar alignment studies can be repeated for other organisms and fixed-variable residue templates for their nanobodies can be obtained.

**Fig. 1.**
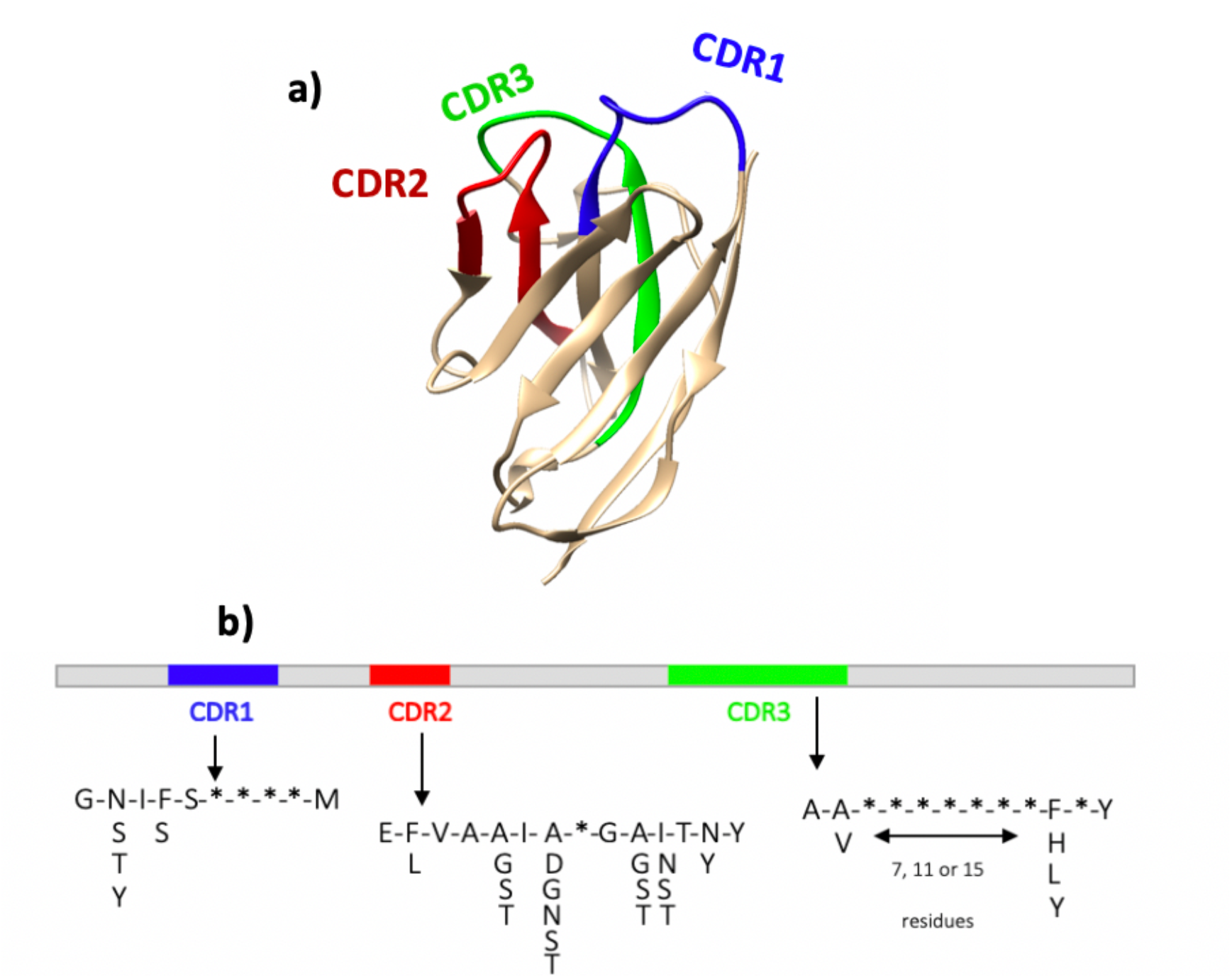
a) Cartoon representation of a nanobody. CDR1, 2 and 3 are colored in blue, red and green respectively. b) Framework of fixed, variable and highly variable amino acid motifs of each CDR are indicated, amino acids indicated with * are the highly variable ones. Residue positions with more than one type of amino acid, which are the variable regions for the Llama glama nanobodies, and single residue positions are also shown.

Computational screening methods contributed greatly to the optimization and design process. There are several approaches available. The common protocol is to start the screening procedure from a known potential binder nanobody retrieved from libraries by phage display or by other selection protocols; bacterial display, yeast display, ribosome display or intracellular 2 hybrid selection, and gradually enhancing the properties by mutating CDR or non-CDR residues iteratively and measure the antigen-nanobody affinity values.

To start the mutation process, the CDR, non-CDR loops and the locations on the CDR loops which allow insertions or deletions must be identified. This can be done by aligning nanobody sequences to known structures belonging to the same species. There are publicly available CDR numbering tools that can be applied to a certain sequence.[5] After the identification of the orientation or the sequence of the nanobody, amino acids can be selected for mutation to increase the binding affinity for the antigen.

Current computational techniques are based on finding the CDR and non-CDR residues that are in contact with the antigen and performing point mutations to enhance the interaction.[6–10] However, there is no certain rule for deciding on which residues to mutate. The optimization strategy proposed in this study is based on finding the hypervariable positions in CDRs by an alignment process and increasing the binding affinity by mutating them iteratively and selecting the residues which are highly frequent for a particular species. This method is tested on several different proteins: human MDM4, Fructose-bisphosphate aldolase from *Trypanosoma congolense* and human lysozyme. PDB codes are 2VYR, 5O0W and 4I0C respectively.

## Method

### Alignment

Sequences and structures of synthetic and natural nanobodies can be found in databases.[11, 12] Nanobody sequences for specific organisms can be downloaded from the databases and aligned by using several multiple alignment tools available. [13–16] The alignments can be visualized by specific softwares [17, 18], the consensus sequence and the conserved or hypervariable positions can be determined and a template can be obtained similar to the llama-derived nanobody framework proposed previously. [4]

Nanobody sequences for *Vicugna pacos* and *Camelus dromedaries* are downloaded from single domain Antibody database (sdAb) [11], and aligned separately by using ClustalW, a multiple sequence alignment tool.[16] Alignments are visualized in Unipro U-gene and hypervariable residues, their evolutionary amino acid frequencies and consensus sequences for CDRs are determined.[17] Residue positions with high gap penalties are not considered, the occurrences of the amino acids in highly variable positions are counted and the frequency of each amino acid is saved to be used further in the energy update step. A consensus sequence is obtained for each alignment and hypervariable positions are labeled by asterisks (*) in the templates. Residues corresponding to these positions will be selected for mutation.

### Optimization Strategy

The strategy that we developed in our study is to start with a bound protein and nanobody complex, determine the hypervariable CDR residues according to the given templates, then perform 20 independent point mutations on variable positions, starting from Alanine up to Valine, by using visual molecular dynamics (VMD).[19] Each mutant structure will be subjected to a minimization, annealing, conventional MD run and another minimization cycle by using NAMD.[20] In this study CHARMM36 force field is used. The interaction energy of the nanobody and the protein for each mutation case is calculated and further updated according to the evolutionary amino acid frequency scheme obtained from the alignment by using an iterative Monte Carlo algorithm which favors higher evolutionary preferences. The end result provides us with the distribution of possible optimum mutation types on hypervariable positions of CDRs. A broad library of optimal nanobodies can be derived and tested for its affinity and selectivity. The affinity of the new nanobodies can be measured by using Steered Molecular Dynamics (SMD)[21], by pulling the nanobody away from the protein and measuring the work required to abolish all of the interactions. These SMD simulations provide the dissociation constant, K_D_ values which are useful in comparing the affinity of the optimized and the wild type nanobodies.

### Molecular Dynamics Min-run cycle

All of the mutant nanobodies in complex with the protein are solvated in a TIP3P water box and counter ions are added to neutralize the system. CHARMM36 force field is used to parametrize the atoms. Time step of simulations are kept as 2 fs. Initially the energy of the system is minimized for 8000 steps and the system is gradually heated up to 310K. The annealing step is followed by an equilibration period at constant temperature of 310 K and constant pressure of 1 bar for 10 000 steps. Finally, the energy of the equilibrated system is minimized for another 5000 steps.

### Interaction Energy Update

The interaction energy includes electrostatic, VdW and non-bonded terms. Majority of the force fields favors the charged residues over the others. According to the alignment studies, stable nanobodies have highly frequent residue types in their hypervariable positions. The interaction energies calculated from the min-run cycle, are updated according to the probability distributions, obtained from the alignment, by using the following algorithm: We assume that the system obeys Boltzmann statistics, and the probabilities are functions of the energy and the temperature of the system. Therefore, the probabilities can be written in the following form;

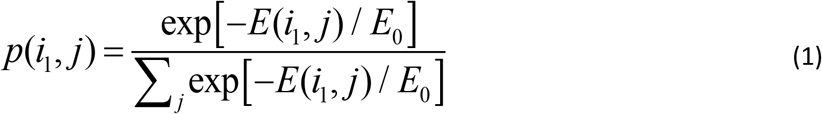

where *E(i_i_,j)* is the interaction energy of each mutant hypervariable position and the antigen obtained from MD min-run cycles, *i*_*1*_ is the hypervariable residue number and *j* is the mutated residue type (Ala, Arg, Asn, …, Val) and *E*_*0*_ scales the temperature. Small values of *E*_*0*_ favor low energy residues for each individual. These priori probabilities are further updated according to the evolutionary distributions obtained from the alignments. The choice of residues according to the a priori probabilities, Eq. 1, is made as follows: For each position, *i*_*1*_, a random number is generated between 0 and 1, and the *j*^*th*^ amino acid mutation for *i*^*th*^ residue is accepted if the random number is between p(i_1_, j) and p(i_1_, j+1). The weights of evolutionary frequencies on the a priori probabilities are introduced as follows: We consider the case where a total of n residues in the nanobody are selected as hypervariable positions. For a given mutated sequence i_1_, where *i*_*1*_ goes from 1 to *n*, we represent the calculated distribution from the MD interaction energies by *C(i)*. The evolutionary distribution for the mutated sequence is *K(i)* where i goes from 1 to *n*. The divergence, *D*_*CD*_, of the distribution *C* from the evolutionary distribution *K* is calculated according to the divergence equation, [22]

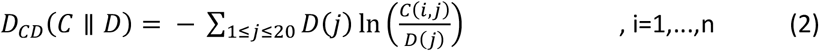

A set of random sequences are generated and the divergences of each random sequence from the evolutionary distribution is measured according to the flowchart shown in Fig.2. Initially the hypervariable positions on the nanobody are detected by comparing the original sequence with the consensus sequence, selected hypervariable positions are mutated to 20 available amino acids and projected to MD min-run cycles. The interaction energies of single mutant and the antigen is measure by using VMD NAMD Energy plugin. Priori probabilities are calculated from the MD interaction energies and these energies are further updated by evolutionary probabilities for each amino acid type for a given species. A library of mutant nanobodies which have better binding properties to the antigen are generated as a result of this process. This library is further filtered by measuring the divergence of residue mutations selected for hypervariable positions from the evolutionary probabilities. The sequences that have lower divergence from the evolutionary distribution are kept and a mutation library is generated with the proposed probability distributions for each hypervariable position. Mutations are selected from the proposed probability distributions and the affinities are calculated by the steered molecular dynamics method.

**Fig 2.**
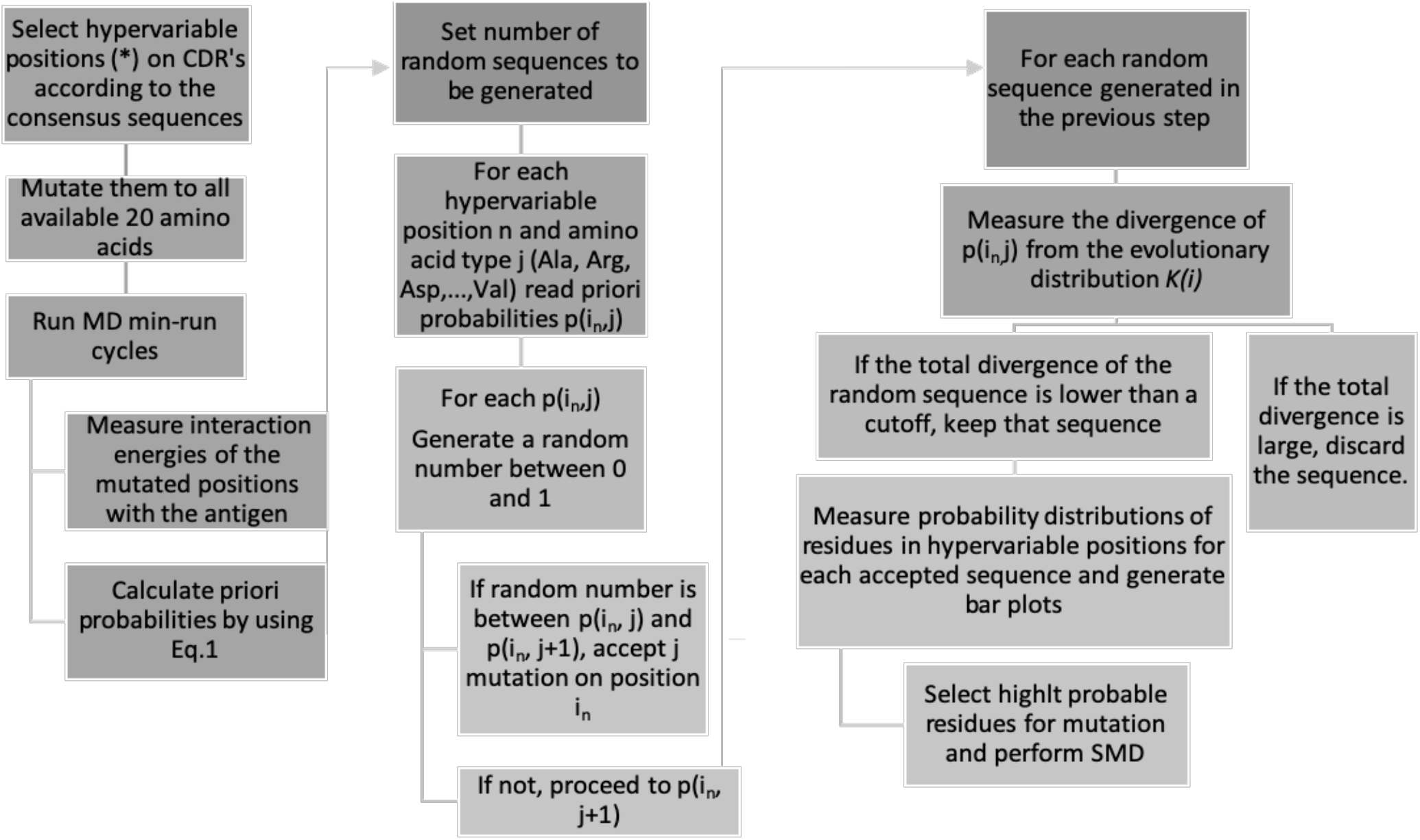
The flowchart of the optimization method.

### Steered Molecular Dynamics

In SMD, the binding affinity is correlated with the rupture force, where the ligand is totally detached from the protein. It has been shown that nonequilibrium pulling work also correlates well with the experimental results. [23, 24] The proposed mutations are applied to nanobodies and the mutant nanobody is pulled away from the protein by SMD simulations at 310 K.[21]For the SMD simulations the protein is anchored from a residue close to its center of mass but away from the nanobody binding site, the nanobody is pulled away with constant velocity from a residue away from the binding site and close to the free-end, where it is not bound to the protein. The force vectors required for the pulling is obtained from the simulation log file. The force vectors at each timestep are normalized with the pulling direction and the normalized force is plotted against the pulling distance. The cumulative is calculated by summing up the area under the force vs distance curve. The distance at which all the interactions between the nanobody and the protein diminished, rupture distance, is detected from the simulations, and the *K*_*D*_ value is calculated from the cumulative work value corresponding to that distance with the following formula:

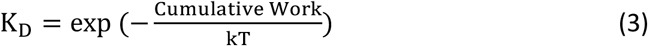

Where the unit of the cumulative work is kj/mol and *kT*, also written as *k*_*B*_*T*, is the product of the Boltzmann constant, *k* and the temperature, *T*. *kT* is 2.58 kJ/mol at 310 K.

10 replicate SMD simulations were performed for each complex and the mean cumulative work values are considered in K_D_ calculations.

## Results

### Alignment results

#### Vicugna pacos

The template for *Vicugna pacos* derived nanobodies are obtained, the consensus sequence for each CDR and the frequency of amino acids at each hypervariable position, shown by asterisks (*), is determined.

#### Camelus dromedaries

The template for *Camelus dromedaries* derived nanobodies are obtained by the same approach. The consensus sequence for each CDR and the frequency of amino acids at each hypervariable position is determined.

The consensus sequence templates shown on Figs 3–5 and Figs 6-8 are obtained from aligning all *Vicugna pacos* and *Camelus dromedaries* derived nanobody sequences available in single domain antibody database. [11] The positions that retain highly variable in the alignments are labeled as the hypervariable positions and the positions that yield in high gap scores are not considered in the alignment. The amino acid frequencies given in Table 1 and 2 are obtained from the amino acid frequencies found in the hypervariable positions within all aligned sequences for *Vicugna pacos* and *Camelus dromedaries* respectively.

**Table 1.**
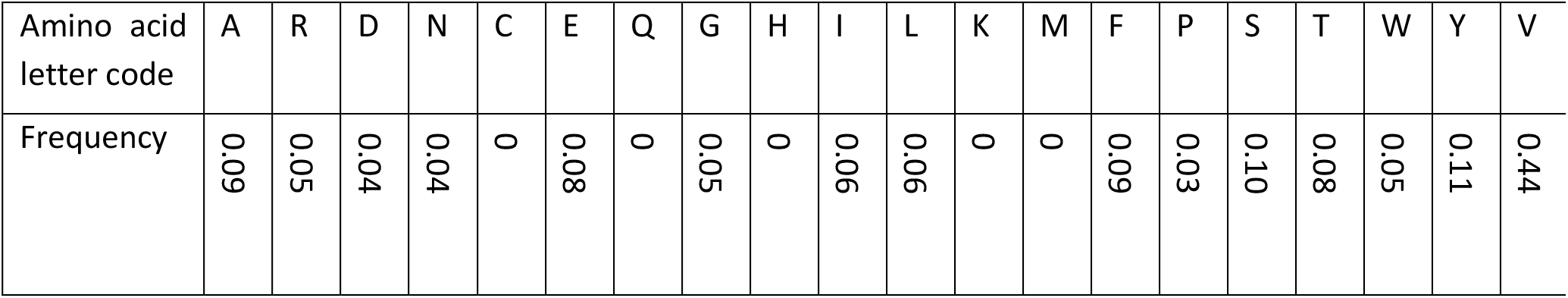
Amino acid frequencies for hypervariable residue positions shown by asterisks (*) in consensus CDR sequences for for *Vicugna pacos*.

**Table 2.** Amino acid frequencies for hypervariable residue positions shown by asterisks (*) in consensus CDR sequences for *Camelus dromedaries*.

| Amino acid letter code | A | R | D | N | C | E | Q | G | H | I | L | K | M | F | P | S | T | W | Y | V |
| --- | --- | --- | --- | --- | --- | --- | --- | --- | --- | --- | --- | --- | --- | --- | --- | --- | --- | --- | --- | --- |
| Frequency | 0.04 | 0.06 | 0.09 | 0.07 | 0 | 0.04 | 0 | 0.11 | 0 | 0.07 | 0 | 0 | 0 | 0.09 | 0.06 | 0.08 | 0.05 | 0 | 0.15 | 0.05 |

**Fig 3.**
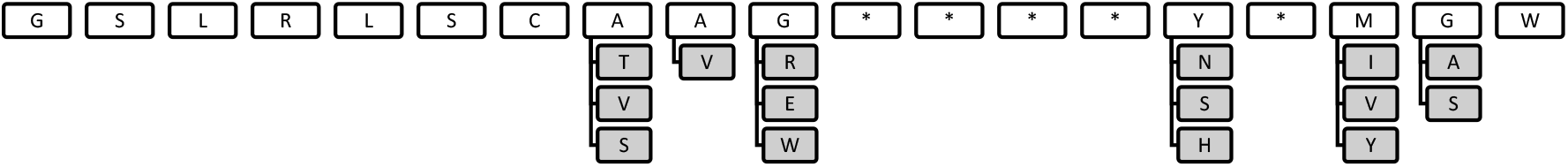
The consensus sequence of CDR1 loop for Vicugna pacos derived nanobodies, the asterisks (*) denote the hypervariable positions in the loops

**Fig 4.**
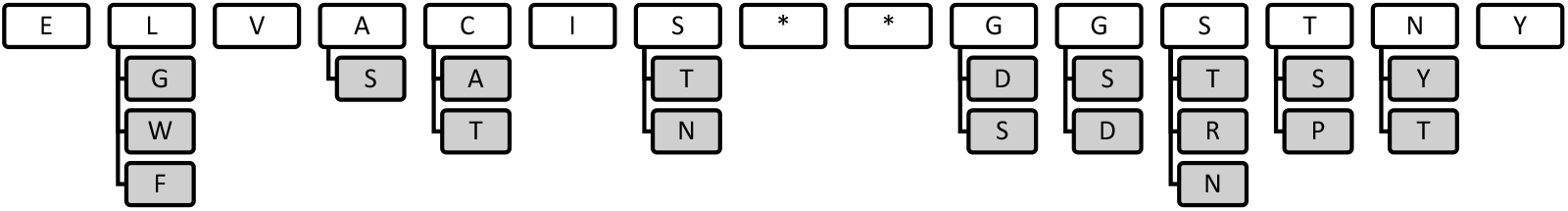
The consensus sequence of CDR2 loop for Vicugna pacos derived nanobodies, the asterisks (*) denote the hypervariable positions in the loops

**Fig 5.**
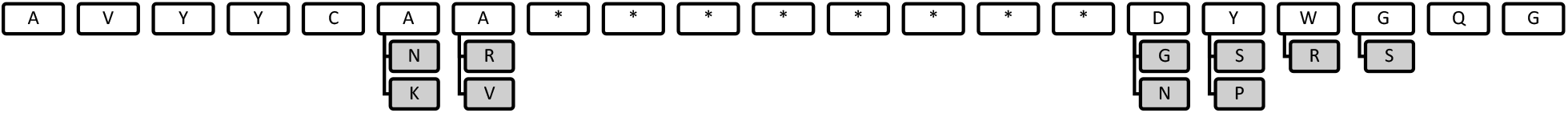
The consensus sequence of CDR3 loop for Vicugna pacos derived nanobodies, the asterisks (*) denote the hypervariable positions in the loops

**Fig 6.**
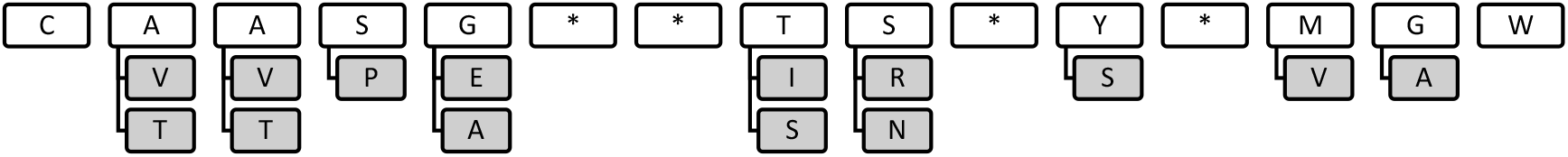
The consensus sequence of CDR1 loop for Camelus dromedaries derived nanobodies, the asterisks (*) denote the hypervariable positions in the loops

#### Optimization results

Nanobody candidates for human MDM4, with PDB id 2vyr, Fructose-bisphosphate aldolase from *Trypanosoma congolense, PDB id 5O0W*, and human lysozyme, PDB id 4I0C were proposed, and the hypervariable mutations are compared with a best binder nanobody to studied targets.

### MDM4

Human MDM4 is a 490 amino acid, key regulatory protein of tumor suppressor p53. It consists of 3 conserved domains. An N terminal for binding p53, a Zinc-finger domain and a C-terminal RING domain. Over expression of them are associated with tumor production and it could be an important regulator in anticancer strategies. MDM4 binds p53 and decreases its activity and stability against degradation. In most of the cancer studies, p53 is inactivated and reactivating p53 is thus an attractive strategy for the treatment. As the result of the recent studies, MDM proteins, negative regulators of p53, seem to be the druggable and controllable oncogenic targets. Several strategies have been proposed to reactivate p53 by suppressing MDM4. These strategies are based on disrupting the p53-MDM4 interaction by either using a small molecule or a peptidic compound. The small molecule antagonists that disrupts this interaction are nutlin and WK-298. [25, 26] Although these compounds help reactivating p53, they do not elicit all the effects of MDM overexpression. Peptidic antagonists have also been developed. Although they have larger interaction surfaces, most of them are unstable in vivo. Another approach is to use mini proteins, nanobodies, to antagonize the interaction between p53 and MDM4. In a study of selecting MDM4 specific single domain antibody, a successful candidate nanobody was found to stabilize and inhibit p53-MDM4 interaction with a dissociation constant of 44 nM. This synthetic construct, VH9, is known to be the best binder for MDM4.[27]

Our aim is to use our optimization strategy to detect the wild type amino acids of the hypervariable positions and propose an amino acid probability distribution for them which will result in similar binding affinity and selectivity with VH9.

Crystal structure for MDM4-VH9 with the PDB id 2VYR is used. VH9 is a synthetic construct, thus, to determine which consensus sequence to utilize in the interaction energy update step, VH9 sequence is aligned with all consensus sequences for *Llama glama, Vicugna pacos* and *Camelus* dromedaries and the sequence similarity is measured to understand the homology of VH9 nanobody with other species. Percent identity score, which refers to a quantitative homology measure, calculated by multiplying the number of matching amino acids by 100 and dividing it by the length of the aligned sequence. Closely related sequences are expected to yield a higher percent similarity. The percent identity matrix of consensus sequences from other species and VH9 is shown on *Table 3.* VH9 and *Llama glama* alignment gives a higher percent similarity score, therefore VH9 is more homologous to *Llama glama* and its consensus sequence can be used to update MD interaction energies of VH9. The template shown on *Fig1.b* for *Llama glama*, obtained from a previous study, is used to update the MD energies.[4] The hypervariable positions were detected according to the *Llama glama* template and the optimization procedure is applied. According to the template, highly variable residue positions for each CDR are residues number 30, 31, 32 and 33 for CDR1, 53 for CDR2 and 94, 95, 96, 97, 98, 99, 100, 101, 102 and 104 for CDR3.

**Table 3.** Sequence identity matrix of nanobody consensus sequences and VH9, measured by Clustal12.1

|  | VH9 | *Vicugna pacos* | *Llama glama* | *Camelus dromaderies* |
| --- | --- | --- | --- | --- |
| VH9 | 100.0 | 62.3 | **69.3** | 66.9 |
| *Vicugna pacos* | 62.3 | 100.0 | 70.8 | 66.4 |
| *Llama glama* | 69.3 | 70.8 | 100.0 | 69.3 |
| *Camelus dromaderies* | 66.9 | 66.4 | 69.3 | 100.0 |

Hypervariable residue mutations with high probability are selected from *Fig 9.* and a mutant nanobody sequence is generated. Selected hypervariable residue mutations and WT residue types are compared in Table 4. The affinity of the mutant nanobody and the wild type VH9 is compared with SMD. A constant pulling speed of 30 Å/ns is applied.

**Table 4.** Residue types at the hypervariable positions for WT VH9 and mutant VH9 optimized for MDM4

|  | 30 | 31 | 32 | 33 | 53 | 94 | 95 | 96 | 97 | 98 | 99 | 100 | 101 | 102 | 104 |
| --- | --- | --- | --- | --- | --- | --- | --- | --- | --- | --- | --- | --- | --- | --- | --- |
| **WT** | GLU | GLU | TYR | ALA | ALA | TYR | TYR | CYS | ALA | LYS | PRO | TRP | TYR | PRO | MET |
| **Mutant** | GLU | TYR | THR | ARG | HIS | TYR | SER | GLY | GLY | TYR | ARG | TYR | ARG | GLY | ASP |

**Fig 7.**
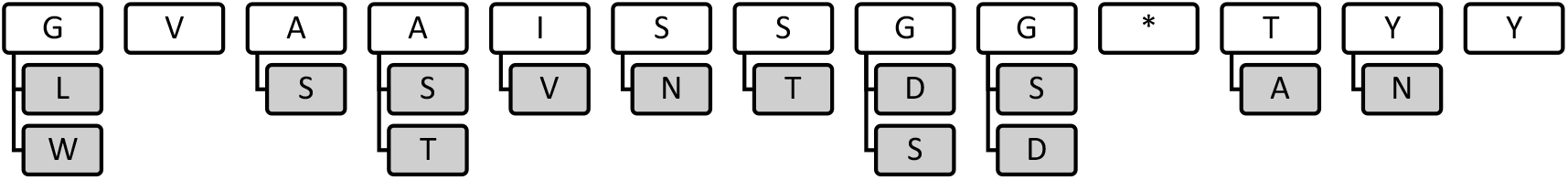
The consensus sequence of CDR2 loop for Camelus dromedaries derived nanobodies, the asterisks (*) denote the hypervariable positions in the loops

**Fig 8.**
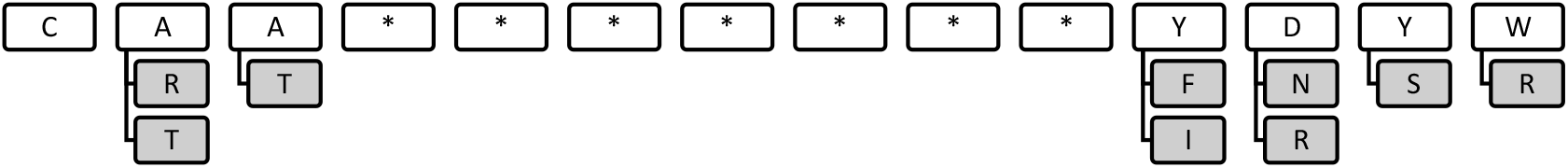
The consensus sequence of CDR3 loop for Camelus dromedaries derived nanobodies, the asterisks (*) denote the hypervariable positions in the loops

**Fig 9.**
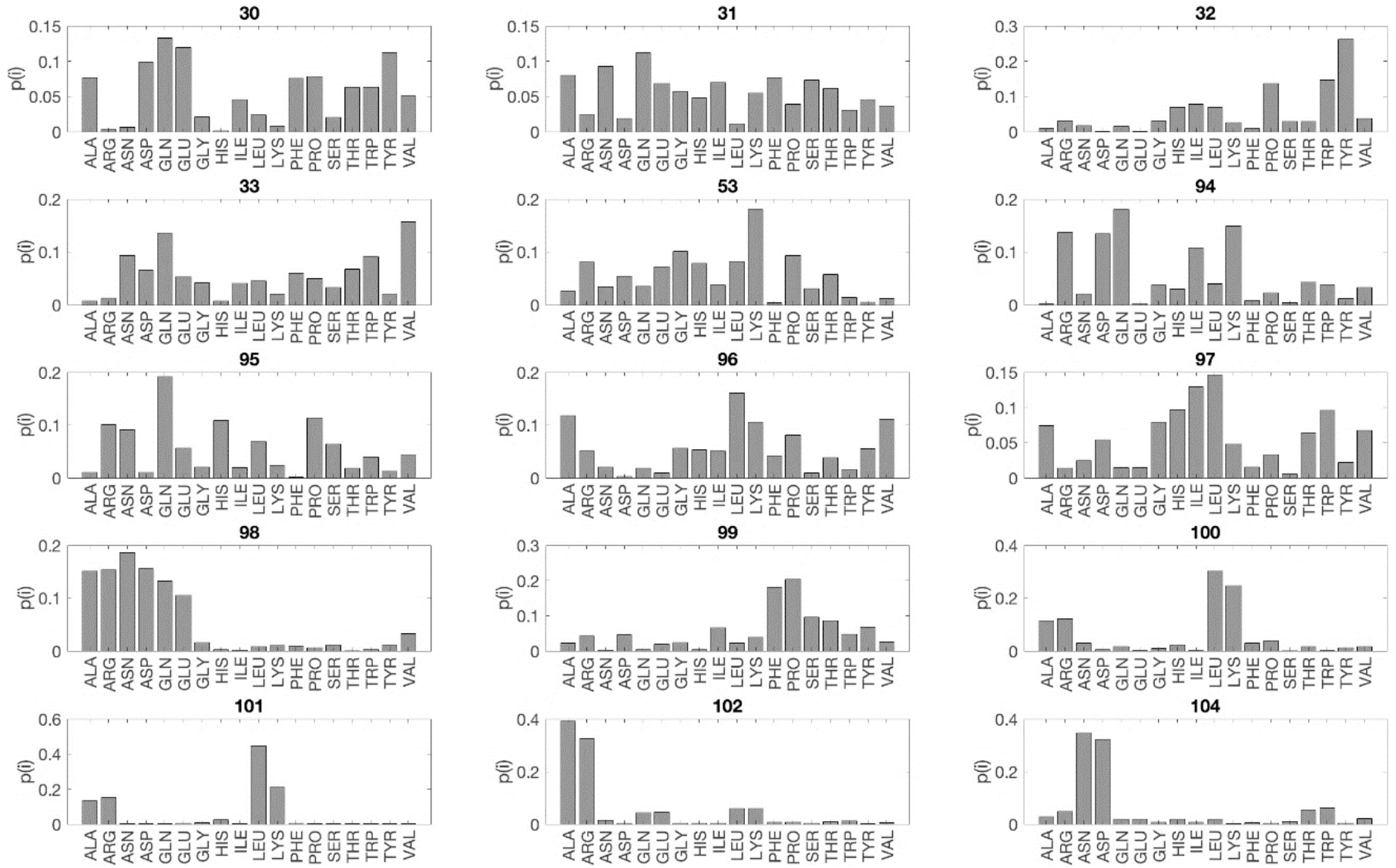
The probability distribution of the optimum residue mutation types for selected hypervariable positions on VH9 in complex with MDM4 obtained by the optimization method described in the flowchart in Fig 2.

Affinities are calculated from SMD plots given in Fig 10. The rupture distance is 10.34 Å for WT VH9 and 11.83 Å for mutant VH9. Force values are obtained from the simulation log files and the cumulative work values are measured by calculating the area under the force vs distance plots. The K_D_ values are calculated from the cumulative work values corresponding to rupture distances. Calculated K_D_ values are 18.12 nM and 9.24 nM for WT and mutant VH9 respectively.

**Fig 10.**
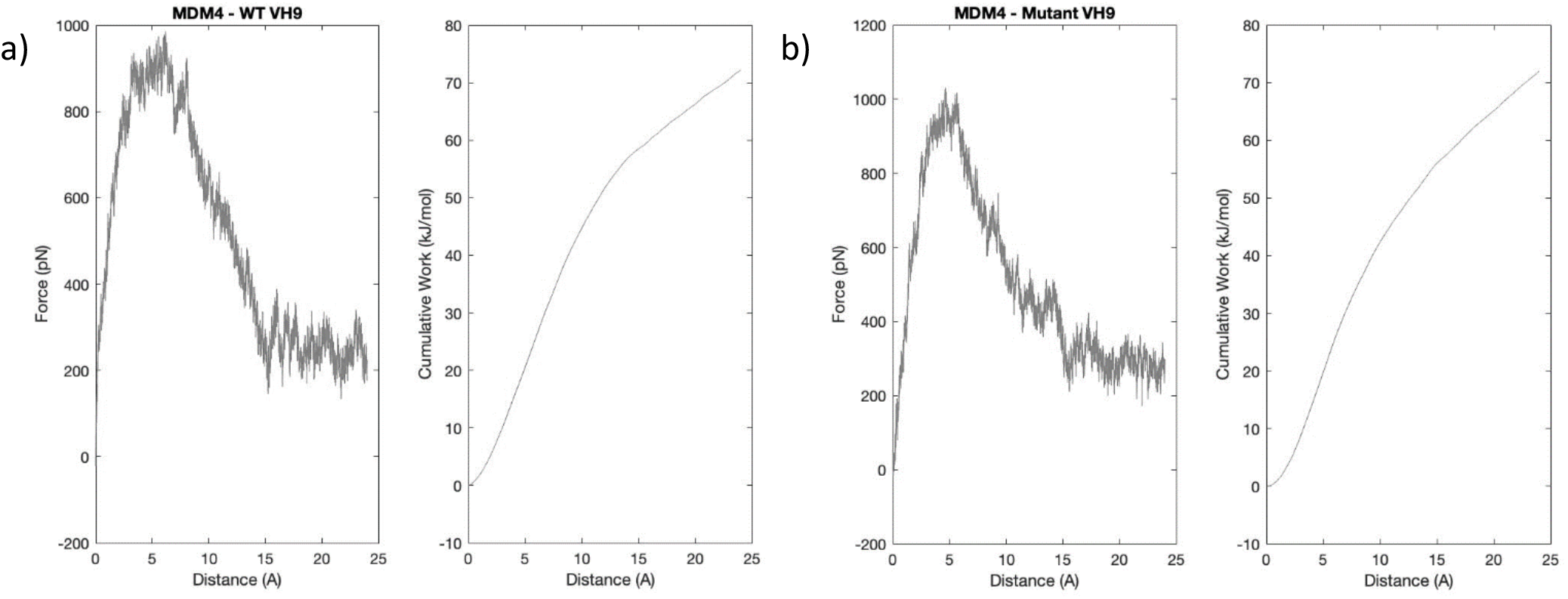
Mean force(pN) vs distance(Å) and mean cumulative work(kJ/mol) vs distance(Å) plots for a) WT VH9 and b) mutant VH9

### Fructose-bisphosphate aldolase from Trypanosoma congolense

Trypanosoma genus belongs to a diverse group of parasites, which cause disease in humans and livestock. Infections of this genus leads to Human African Trypanosomosis (HAT) and Animal African Trypanosomosis (AAT). Specifically, *Trypanosoma congolense* is responsible for the infections and it also leads to major economic losses. A proper diagnostic tool is required in order to detect the infection in animals and proceed with the selective treatment. Recent studies show that nanobodies can be used as a diagnostic tool to recognize *Trypanosoma congolense* fructose-1,6-bisphosphate aldolase (TcoALD). It is a well conserved glycolytic enzyme among Trypanosoma genus. The affinity of the nanobody, Nb474, determined for enzyme detection has a dissociation affinity constant of 73.83 pM.[28]

The aim is to use this optimization strategy to propose similar amino acids of the WT Nb474 hypervariable positions which will yield in the similar binding affinity and the selectivity with Nb474. Crystal structure for TcoALD-Nb474 with the PDB ID 5O0W is used. The template for *Vicugna pacos* is used to update the MD energies, since it is an alpaca derived Nb. The hypervariable positions were detected according to the template and the optimization procedure is applied. According to the template, highly variable residue positions for each CDR are residues number 28, 31, 32 for CDR1, 53 for CDR2 and 103, 104, 105, 106, 108, 110, 115, 125 and 126 for CDR3.

Hypervariable residue mutations which have the highest probabilities given in Fig. 11 are selected to replace WT residues at each hypervariable position. Selected hypervariable residue mutations and WT residue types are compared in Table 5. The affinity of the mutant nanobody and the wild type Nb474 is compared with SMD. A constant pulling speed of 30 Å/ns is applied.

**Table 5.** Residue types at the hypervariable positions for WT Nb474 and mutant Nb474 optimized for TcoALD

|  | 28 | 31 | 32 | 53 | 103 | 104 | 105 | 106 | 108 | 110 | 115 | 125 | 126 |
| --- | --- | --- | --- | --- | --- | --- | --- | --- | --- | --- | --- | --- | --- |
| **WT** | ALA | TYR | TYR | ARG | ASP | THR | THR | ASP | TYR | SER | TYR | ASP | TYR |
| **Mutant** | ARG | GLU | ALA | HIS | ASN | ASP | LEU | GLN | LEU | LYS | TYR | ASP | GLN |

**Fig 11.**
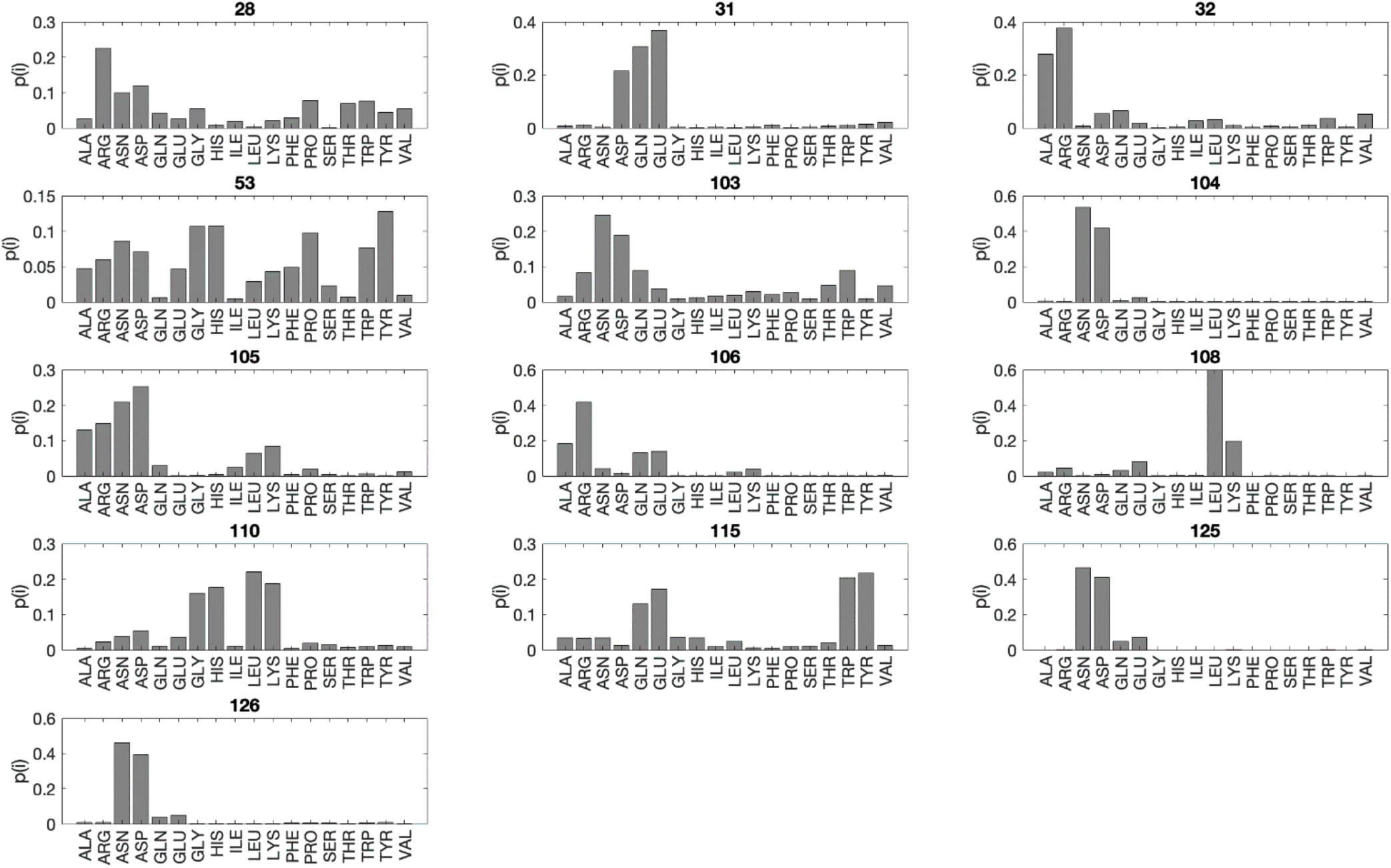
The probability distribution of optimum residue mutation types of selected hypervariable positions of Nb474 in complex with TcAldolase obtained by the optimization method described in the flowchart in Fig 2.

Affinities are calculated from SMD plots given in Fig 12. The rupture distance is 11.09 Å for WT Nb474 and 12.07 Å for mutant Nb474. Force values are obtained from the simulation log files and the cumulative work values are measured by calculating the area under the force vs distance plots. The K_D_ values are calculated from the cumulative work values corresponding to rupture distances. Calculated K_D_ values are 31.22 pM and 634.5 pM fot WT and mutant Nb474 respectively.

**Fig 12.**
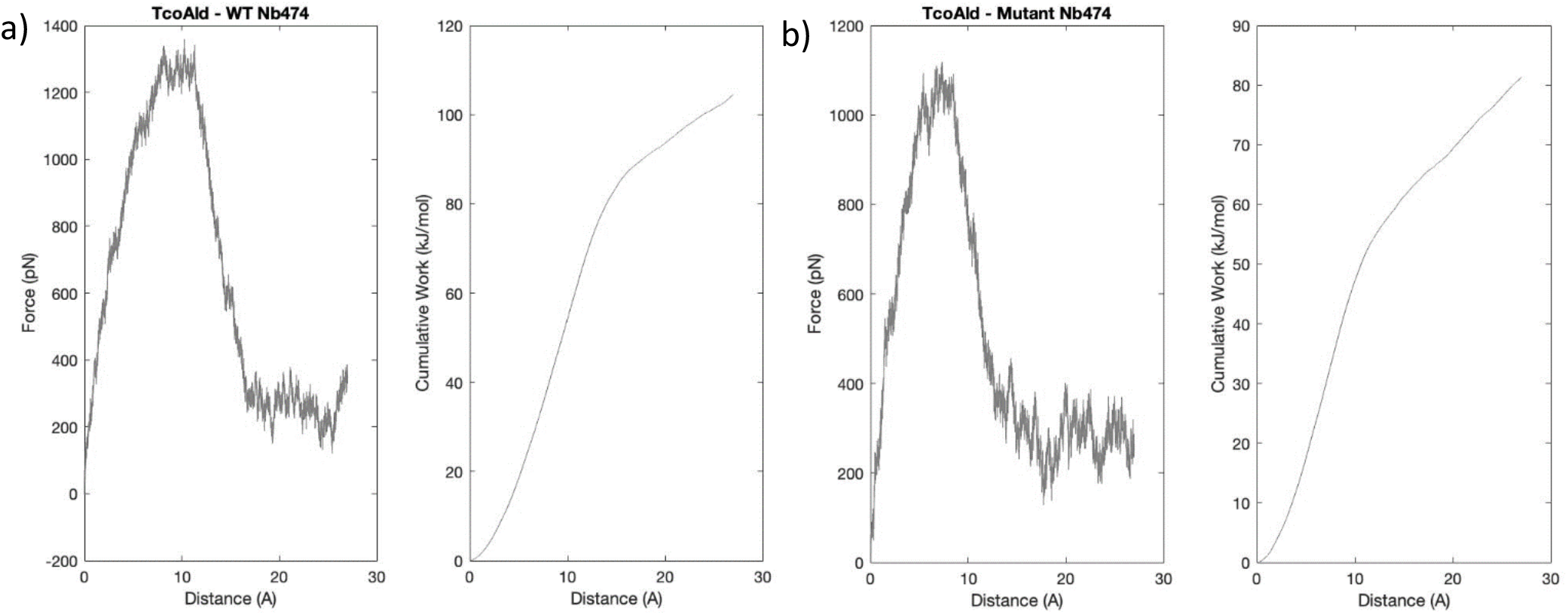
Mean force(pN) vs distance(Å) and mean cumulative work(kJ/mol) vs distance(Å) plots for a) WT Nb474 and b) mutant Nb474

### Human Lysozyme

Human lysozyme (HuL) belongs to the c-type class lysozymes. It is a 130 amino acid protein, capable of both hydrolysis and trans-glycosylation, plays a vital role in host defense. Several variants of this protein were found to be related with a familial systemic non-neuropathic amyloidosis. It has been revealed that several nanobodies are available that prevent the formation of pathogenic aggregates and inhibit fibril formation by the amyloidogenic variants. These nanobodies, cAbHuL5 and cAbHuL5G, can be used to treat protein misfolding diseases. cAbHuL5 and cAbHuL5G displays an affinity of 460 nM and 310 nM for wild type HuL.[29]

The aim is to use this method to propose similar amino acids of the WT cAbHuL5 hypervariable positions which will yield similar binding affinity and the selectivity with cAbHuL5.

Crystal structure for HuL-cAbHuL5 with the PDB ID 4I0C is used. The template for *Camelus dromedaries* is used to update the MD energies. The hypervariable positions were detected according to the template and the optimization procedure is applied. According to the template, highly variable residue positions for each CDR are residues number 27, 28, 29, 30, 31 for CDR1, 55, 56 for CDR2 and 97, 98, 101, 110, 111, 112, 113 and 115 for CDR3.

Optimum hypervariable position mutations are selected from the probability distributions given in Fig 13. A mutant nanobody is generated and the affinity of the mutant nanobody and the wild type CaBHuL5 is compared with SMD. Selected hypervariable residue mutations and WT residue types are compared in Table 6. A constant pulling speed of 30 Å/ns is applied.

**Table 6.** Residue types at the hypervariable positions for WT cAbHuL5 and mutant cAbHuL5 optimized for HuL

|  | 27 | 28 | 29 | 30 | 31 | 55 | 56 | 97 | 98 | 101 | 110 | 111 | 112 | 113 | 115 |
| --- | --- | --- | --- | --- | --- | --- | --- | --- | --- | --- | --- | --- | --- | --- | --- |
| **WT** | LEU | SER | THR | THR | VAL | PHE | PRO | LYS | THR | PHE | SER | ARG | ALA | TYR | HIS |
| **Mutant** | LEU | ASN | ASP | GLU | GLN | ASP | GLN | ALA | LEU | LEU | TRP | SER | LYS | TRP | VAL |

**Fig 13.**
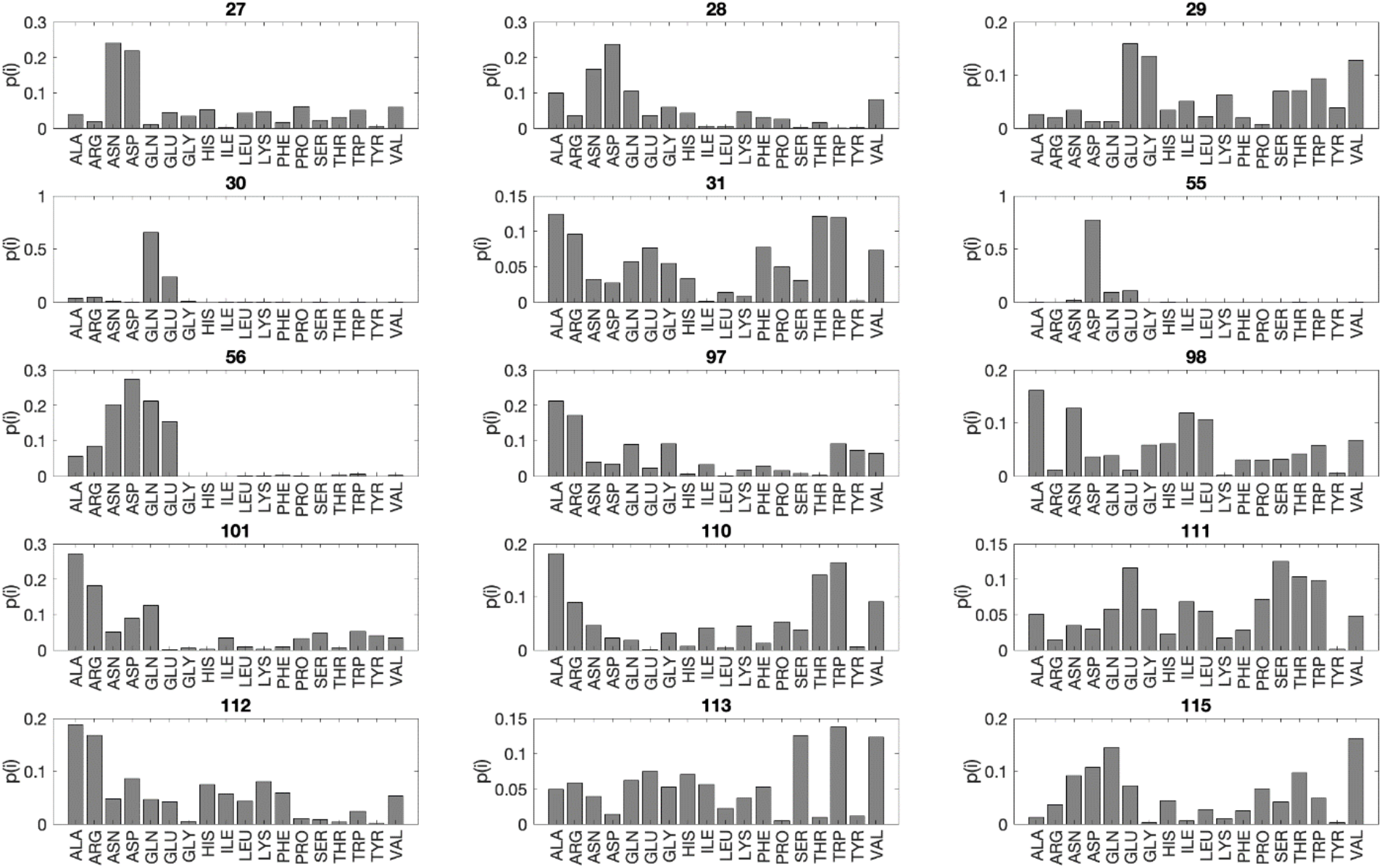
The probability distribution of optimum residue mutation types of selected hypervariable positions of cAbHuL5 in complex with HuL obtained by the optimization method described in the flowchart in Fig 2.

Affinities are calculated from SMD plots given in Fig 14. The rupture distance is 11.01 Å for WT cAbHuL5 and 12.11 Å for mutant cAbHuL5. Force values are obtained from the simulation log files and the cumulative work values are measured by calculating the area under the force vs distance plots. The K_D_ values are calculated from the cumulative work values corresponding to rupture distances. Calculated K_D_ values are 0.15 nM and 1.48 pM fot WT and mutant cAbHuL5 respectively.

**Fig 14.**
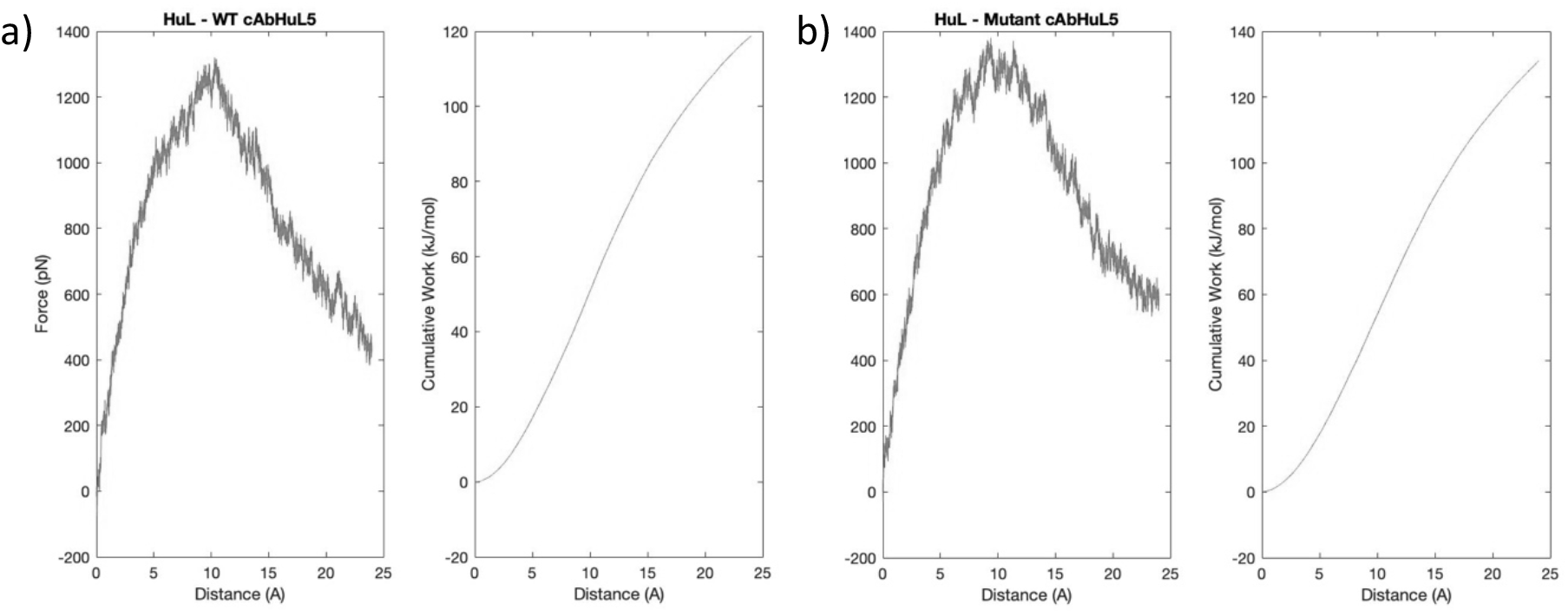
Mean force(pN) vs distance(Å) and mean cumulative work(kJ/mol) vs distance(Å) plots for a) WT cAbHuL5 and b) mutant cAbHuL5

## Discussion

In this article we present a method to find better binding nanobodies for protein targets. Binding of known protein-nanobody complexes are used as benchmark studies. Evolutionarily variant residues on nanobodies are determined and nanobodies with similar affinities to the best binders are generated. The K_D_ values for wild type nanobodies are replicated with 10 replicates of Steered Molecular Dynamics simulations and the affinity of mutated nanobodies are compared with the results of experimentally detected values.

### MDM4-VH9

VH9 is the best binding nanobody to human MDM4 N-terminal domain synthesized so far. The experimentally determined dissociation constant of VH9 bound to human MDM4 N-terminal domain is 44 nM. [27] The dissociation constant we detected in our SMD simulations is 18.12 nM, which is in the same nM range as the experimentally determined value. Hypervariable amino acids in the wild type VH9 falls into the highly probable amino acid scheme that is proposed by this optimization method, shown in Fig.2. The hypervariable amino acids of VH9 are mutated according to the probability distribution scheme and the dissociation constant of the mutant VH9 was measured as 9.24 nM.

The contact points shown in a and b panels of Fig.15 show that this optimization strategy helps us to finding nanobodies that preserve the wild type interactions. The contact points between the antigen and the mutant nanobody is increased. The arrows on Fig15 b. show the enhanced interacting sites between MDM4 and the CDR loops of the nanobody upon mutation. The measured K_D_ values for the mutant nanobody shows that the optimization method is promising and nanobodies with similar affinities to experimentally detected best binding nanobodies can be detected. The interaction of non-CDR residues and MDM4 in mutant VH9 is improved. The binding surface area, calculated by using Cocomaps, of the nanobody with MDM4 is increased from 792.2 Å^2^ to 893.55 Å^2^ in mutant structure. [30] The affinity of both nanobodies for MDM4 is in nM range. Pairwise interacting residues of the WT and mutant nanobody are compared on FigS1. The mutations on CDR2 loop increased the interaction between the antigen and the nanobody. Increased interactions between them caused an increase in the binding affinity of the nanobody.

**Fig 15.**
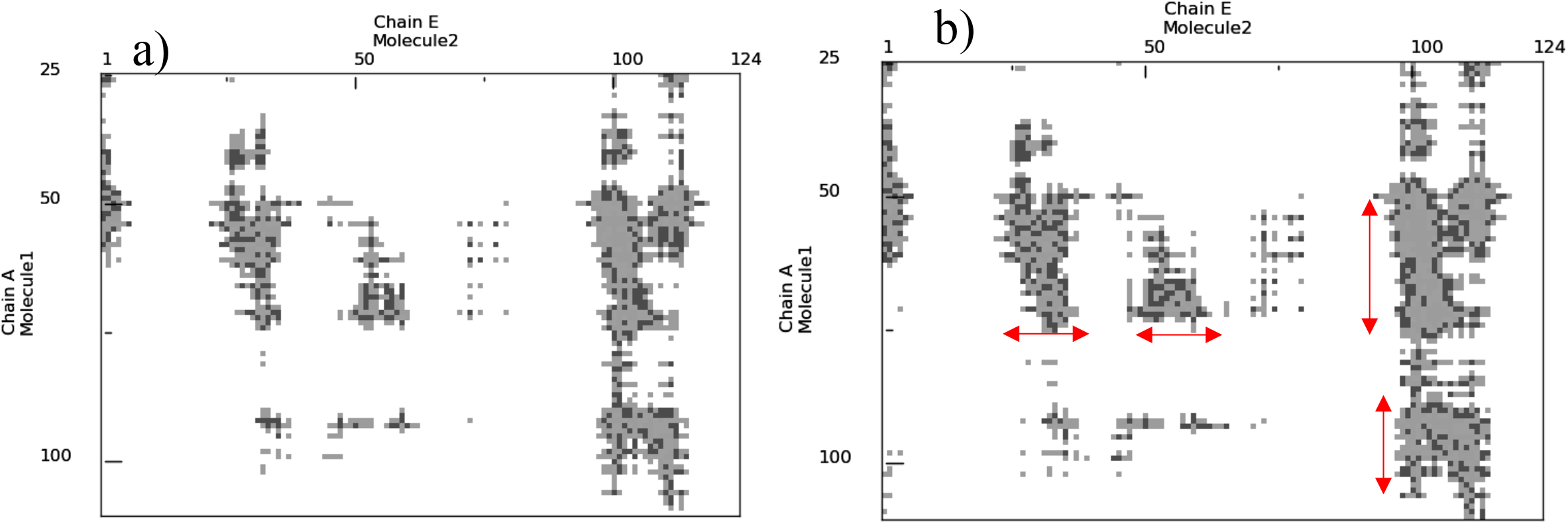
Contact maps of a) MDM4 and WT VH9 and b) MDM4 and mutated VH9. Chain A denotes the MDM4 and chain E denotes the nanobody in both maps, generated using Cocomaps.The red arrows on panel b highlights the increased contacts of the MDM4-VH9 complex upon optimization.

### TcoAld-Nb474

A diagnostic assay, employing the smallest antigen binding Nb474, was prepared to detect *T*. *congolense* infections in animals. This nanobody specifically recognizes *T*. *congolense* fructose-1,6-bisphosphate aldolase. The affinity of this specific interaction is experimentally found as 73.83 pM, meaning a very tight binding, which we determined as 31.22 pM in our SMD simulations.[28] The mutant Nb474 generated by mutating hypervariable residues according to the probability distribution scheme proposed for TcoAld - Nb474 template, shown in Fig.4. The affinity of the mutant Nb474 for TcoAld is measured as 634.5 pM. Both WT and the mutant Nb474 have affinities in the same magnitude of order.

The contact maps in Fig.16 and SMD plots in Fig. 12 show that; similar binding patterns and affinities can be achieved by using this optimization strategy with experimentally detected nanobodies. The contacting pairs in a and b panels of Fig16. are indistinguishable, with some slight increased contacts between the antigen and CDR2 loop of the nanobody, although 13 mutations were applied to the nanobody. The TcAldolase-nanobody interface area is calculated by using Cocomaps, an online tool to analyze protein interaction interfaces [30], as 820.75 Å^2^ and 915.55 Å^2^ for WT and mutant cases respectively. Both affinities of mutant and WT nanobodies are in pM range and interactions in WT Nb474 are preserved in the mutant case and the binding surface area is increased. Residue interactions shown on FigS2. Indicated that our optimization strategy enhanced CDR1 interactions with the antigen.

**Fig 16.**
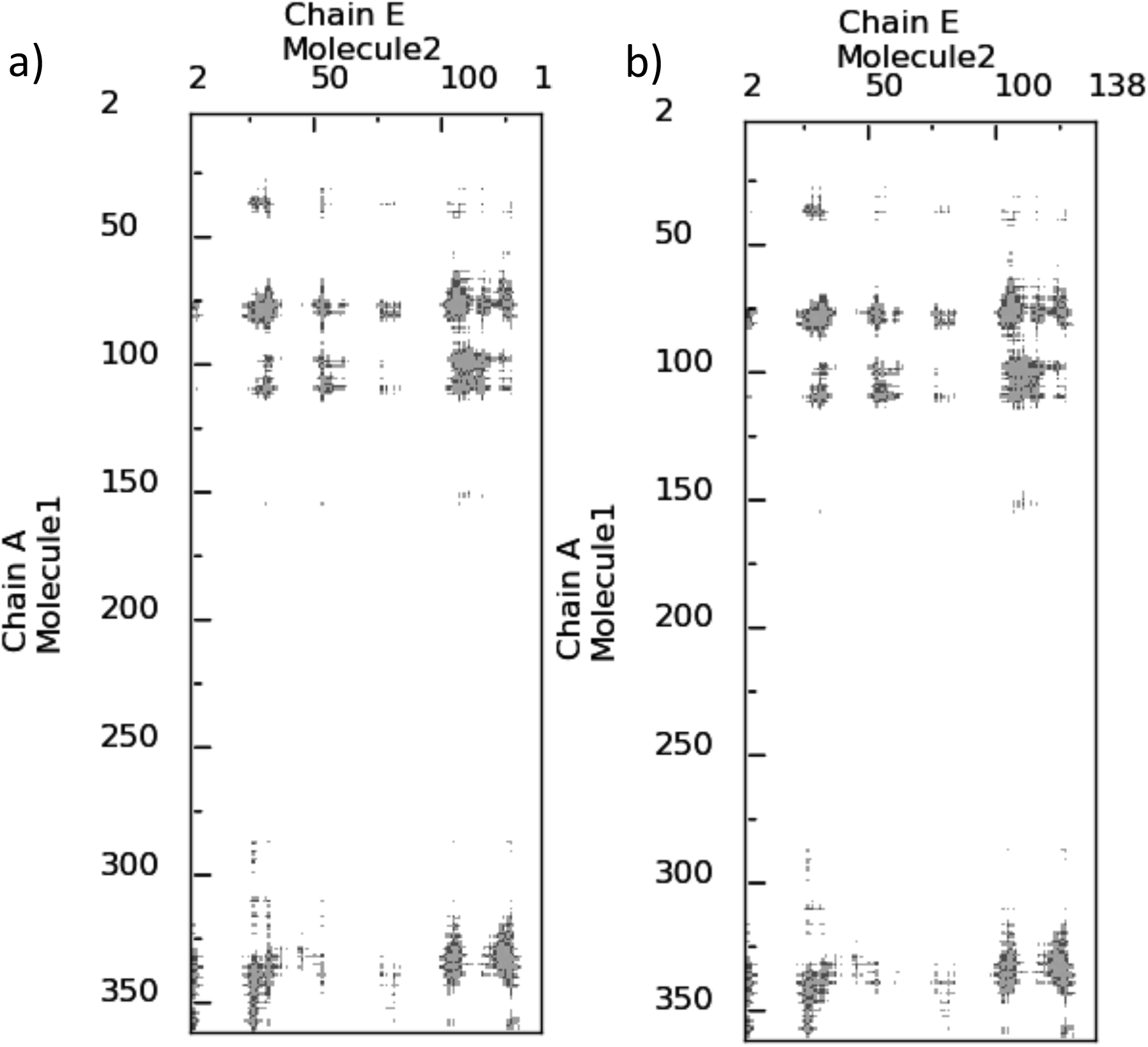
Contact maps of a) TcAldolase and WT Nb474 and b) TcAldolase and mutated Nb474. Chain A denotes the TcAldolase and chain E denotes the nanobody in both map, generated using Cocomaps.

**Fig 17.**
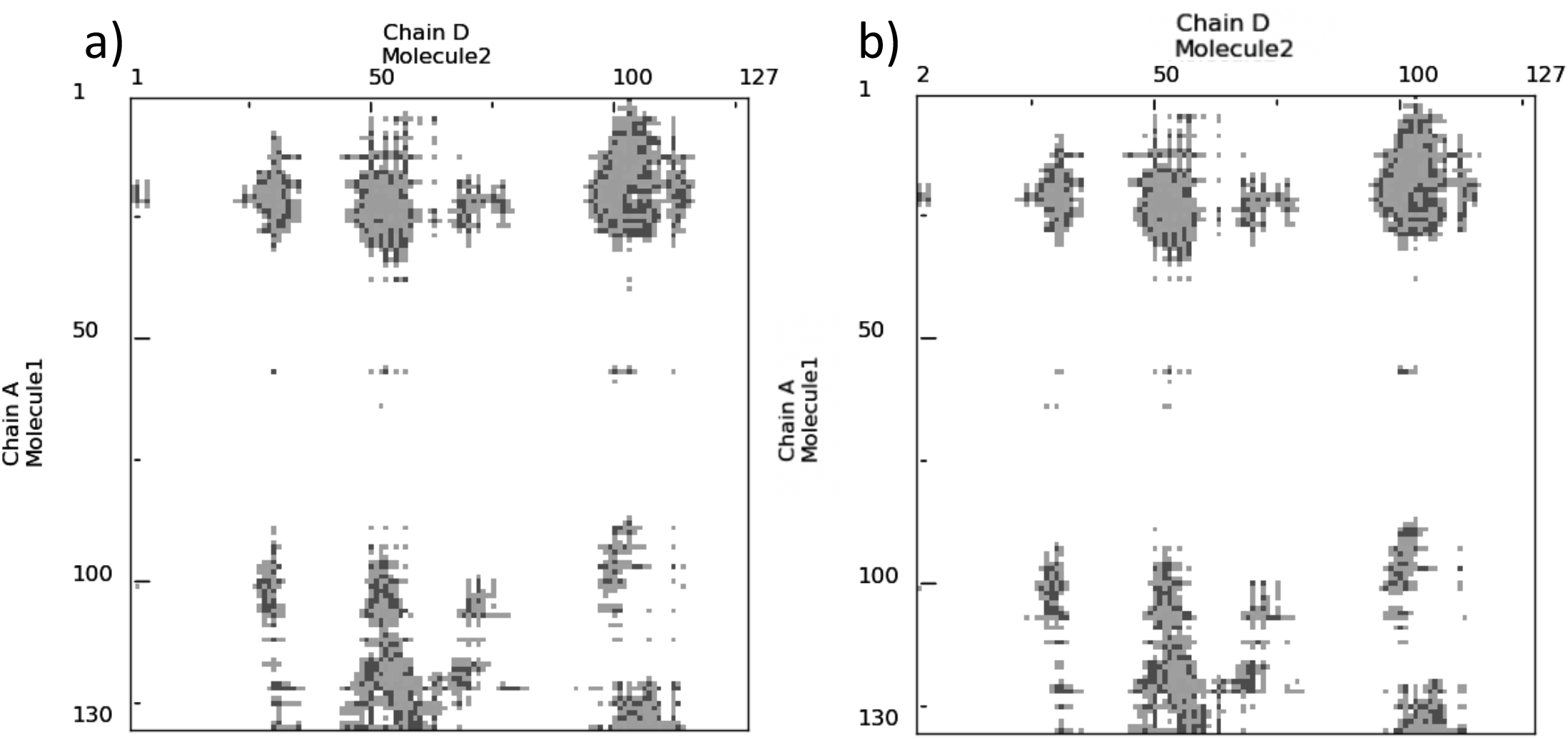
Contact maps of a) HuLand WT cAbHuL5 and b) HuL and mutated cAbHuL5. Chain A denotes the HuL and chain D denotes the nanobody in both map, generated using Cocomaps.

#### HuL-cAbHuL5

The reported nanobody inhibits fibril formation of Human Lysozymess by preventing the unfolding and structural reorganization of the α-domain. The studies provide evidence that nanobodies can be used as therapeutic reagents for protein misfolding diseases. The suggested nanobody has an affinity of 460 nM towards HuL, which is obtained as 0.15 nM in SMD simulations in this study.[29] The proposed mutant according to the optimization method shows an affinity in low pM range, 1.48 pM.

The nanobody generated by mutating its hypervariable CDR residues have higher affinity towards HuL. Similar contact sites were achieved with higher affinity upon mutating hypervariable residues. The binding surface are increased from 662.95 Å^2^ to 691.6 Å^2^ upon mutation. We can conclude that tighter binding nanobodies can be proposed by this optimization method. Individual residue interactions for WT and mutant cAbHuL5 are shown in FigS3. WT cAbHuL5-antigen residue interactions are preserved when hypervariable residues on CDR loops are mutated and the mutations yielded tighter binding.

Therapeutic nanobodies are currently emerging, powerful biological drugs used for treatment and diagnostic purposes. Their small size, increased stability and solubility and large binding regions make them great therapeutic reagent candidates. Experimentally screening the best binder by phage display based methods is time-consuming and computational tools, such as molecular dynamics and steered molecular dynamics can be used to design a nanobody for a specific target and characterize the binding properties. In this study, an optimization method is proposed by selecting the evolutionarily variable residues on nanobodies to design conformationally selective nanobodies and finding amino acid types for hypervariable residues that improve nanobody-protein interaction by applying point mutations on them and determining their contribution in binding energy and binding. Affinities of the mutant and WT nanobodies can be compared by using steered molecular dynamics. The unbinding direction can be optimized by comparing the SMD results with the actual experimental results.

This study reveals that mutation on hypervariable CDR residues improve the binding affinity of nanobodies and can detect experimentally proven best binders. Currently proposed method can be extended for wider applications and many tighter binding nanobodies can be designed for specific targets within nM-pM affinity range. We conclude that computationally optimizing antigen-nanobody binding properties with this proposed method will generate a reliable source of nanobody candidates for specific targets and it should be useful in vast variety of immunotherapeutic applications.

