## Supplementary Figures for "ModiBodies: A computational method for modifying nanobodies to improve their antigen binding affinity and specificity"

**Department of Chemical and Biological Engineering, Koc University, Istanbul, Turkey**

### Supplementary Figures

a)

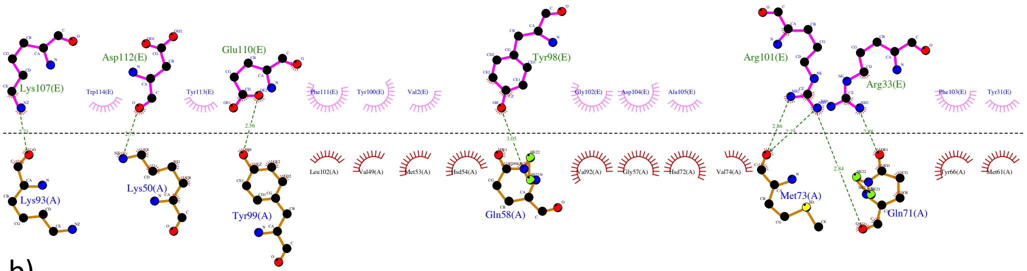


b)

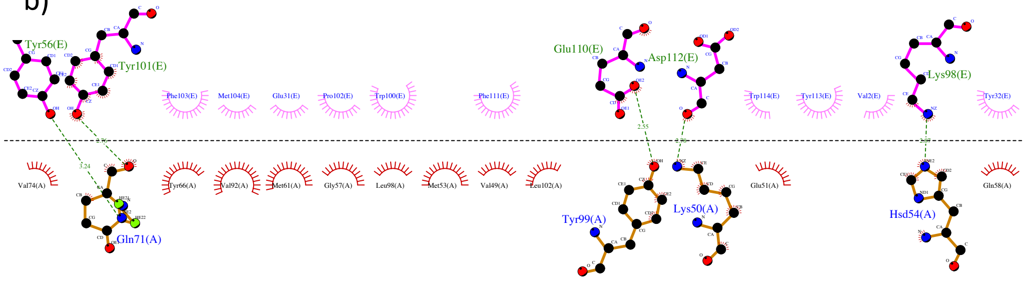


***Fig 1S.*** *Interacting residues of a) MDM4 – WT VH9 and b) MDM4 – Mutant VH9 is shown using Ligplus, where chain A denotes the MDM4 and chain E denotes the nanobody in both figures.*

a)

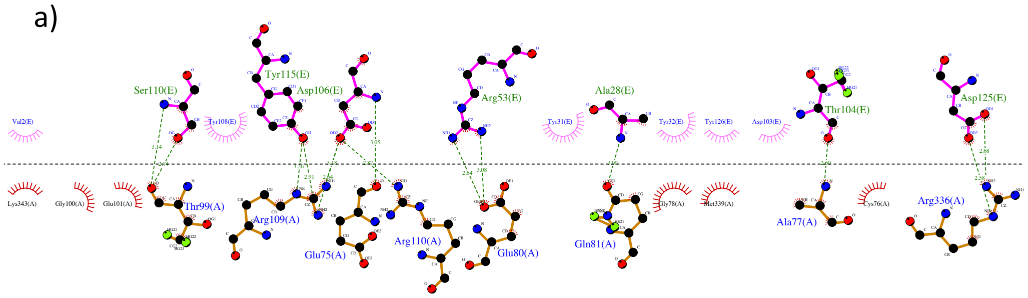


b)

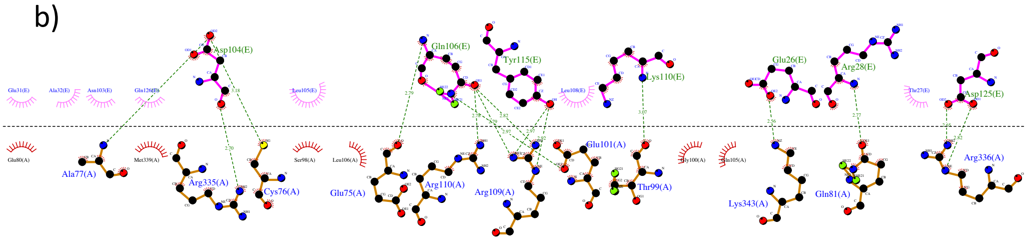


***Fig 2S.*** *Interacting residues of a) TcoAld – WT Nb474 and b) TcoAld – Mutant Nb474 is shown using Ligplus, where chain A denotes the TcoAld and chain E denotes the nanobody in both figures.*

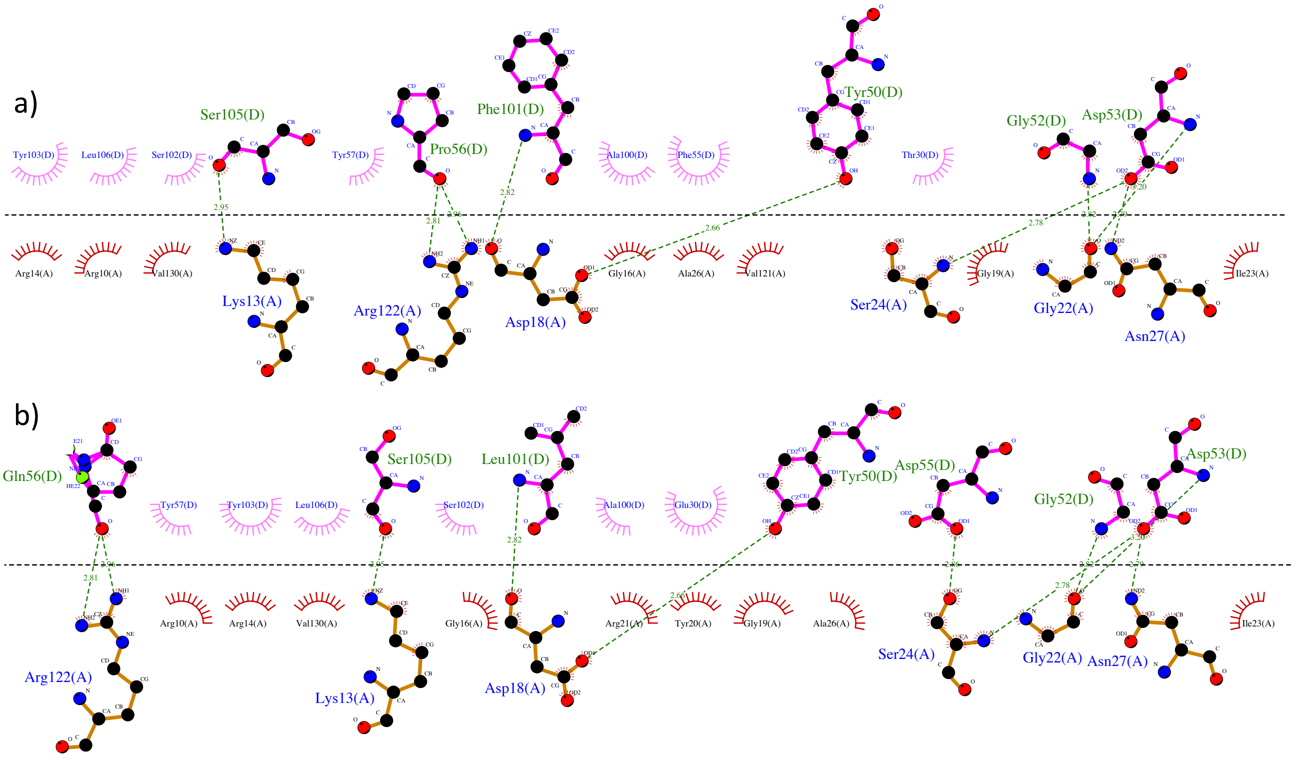


***Fig 3S.*** *Interacting residues of a) HuL – WT* cAbHuL5 *and b) HuL – Mutant* cAbHuL5 *is shown using Ligplus, where chain A denotes the HuL and chain D denotes the nanobody in both figures.*
